# RISC-bound small RNA sequencing provides insights into guide strand selection and siRNA trimming and tailing following insecticidal dsRNA delivery

**DOI:** 10.64898/2026.09.18.748846

**Authors:** Gözde Güney, Paul Plathner, Stefan Scholten, Doga Cedden

## Abstract

RNA interference (RNAi) offers a sequence-specific approach to pest control. In insects, Dicer-2 processes double-stranded RNA (dsRNA) into small interfering RNA (siRNA) duplexes, from which the RNA-induced silencing complex (RISC) retains a guide strand. Only antisense-loaded RISC can mediate cleavage of the target transcript. However, how sequence features shape the RISC-bound siRNA pool in pests remains poorly understood, limiting opportunities for sequence optimization. Here, we profiled RISC-bound siRNAs following injection of 34 insecticidal dsRNAs targeting 11 essential genes in *Tribolium castaneum* larvae. We computationally reconstructed 7,879 siRNA pairs and examined associations between sequence features and strand bias. Differences in GC identity at terminal paired positions 1–5, used as a proxy for local thermodynamic asymmetry, correlated with strand bias, with the strongest correlations at the first two paired positions. ORF targeting and reduced predicted antisense self-folding were also associated with higher antisense fractions. Analysis of non-templated terminal additions revealed predominantly 3′ uridylation, a known signature of small RNA turnover, along with putative 3′ trimming. Among ORF-associated siRNA pairs, sense strands showed higher relative U-tailing abundance, based on 3′-uridylated and putatively trimmed-and-3′-uridylated reads relative to perfect 21-nt reads, than antisense strands. Antisense strands with the least predicted self-folding also showed low relative U-tailing abundance. These observations are consistent with sequence-dependent contributions from both guide-strand selection and differential post-RISC-loading siRNA retention, although a causal link remains to be established. The identified associations provide a basis for testing whether dsRNA sequence optimization can improve pest control efficacy and reduce off-target activity.

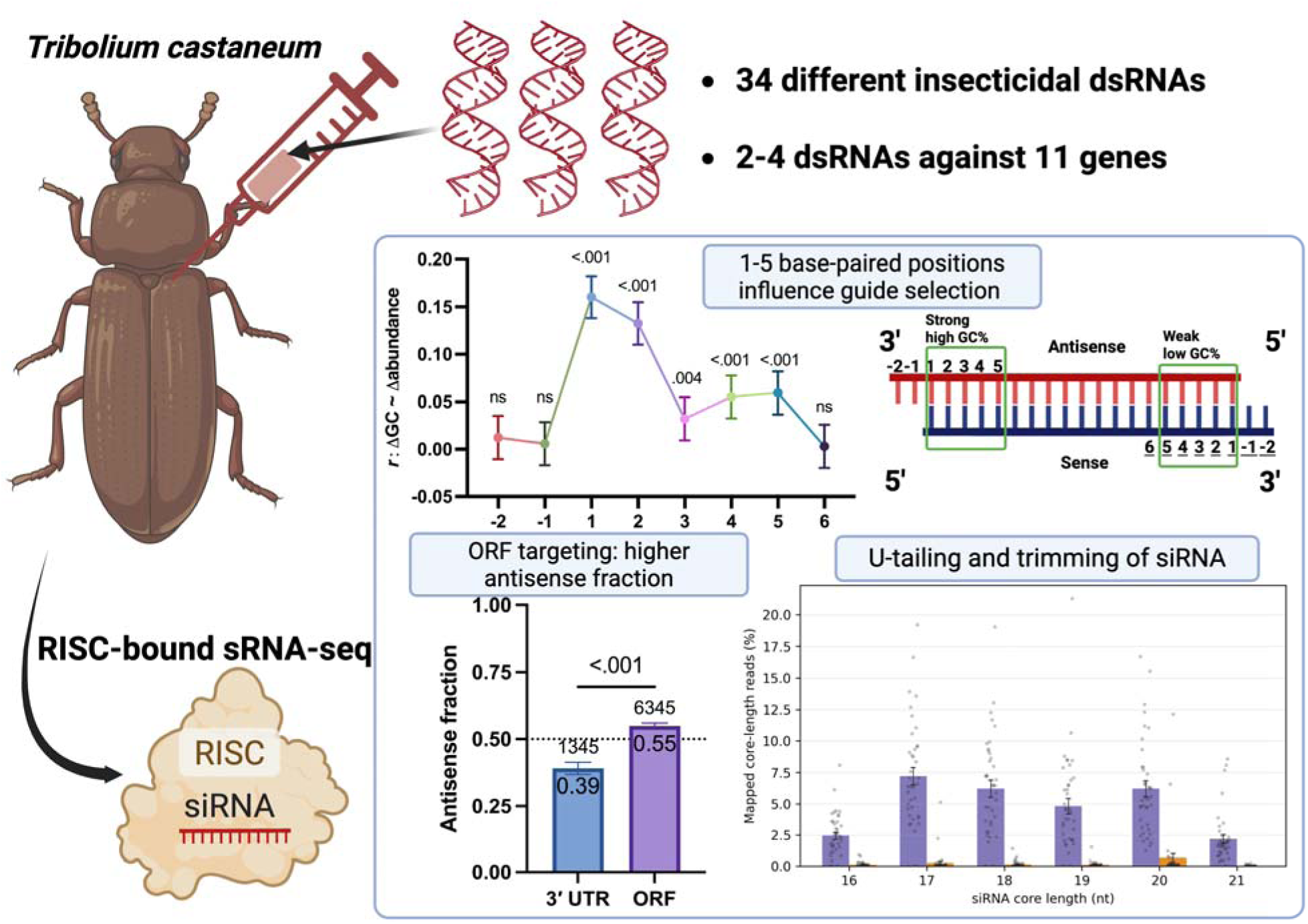

- RISC-bound siRNAs were profiled after delivery of 34 dsRNAs in *T. castaneum*
- Asymmetry in pairing strengths of positions 1-5 correlates with guide strand bias
- ORF targeting and reduced self-folding correlate with higher antisense fractions
- ORF targeting and self-folding sense siRNAs show higher uridylation/trimming

## 1. Introduction

Insect pests cause substantial yield losses in major crops, with annual losses estimated at up to 40% and expected to increase due to climate change (Deutsch et al., 2018). In addition, insects such as mosquitoes transmit pathogens that cause serious human diseases (Farajollahi et al., 2011; van den Berg et al., 2021). Chemical insecticides, including pyrethroids and neonicotinoids, are widely used to control insect pests and disease vectors (Sparks and Nauen, 2015). However, repeated use of insecticides with a limited range of modes of action selects for resistant populations, reducing their effectiveness (Chen et al., 2023; Mallet, 1989; van den Berg et al., 2021). In parallel, broad-spectrum toxicity associated with chemical insecticides poses risks to non-target organisms in the environment, including pollinators (Lundin et al., 2015; Nicholson et al., 2024). These challenges have increased interest in alternative management strategies, including RNA interference (RNAi).

In RNAi-based pest control, double-stranded RNA (dsRNA) complementary to essential pest genes is delivered through genetically engineered crops expressing dsRNA (e.g., SmartStax PRO maize targeting *Diabrotica virgifera virgifera*) or sprayable dsRNA formulations (e.g., Calantha targeting *Leptinotarsa decemlineata*) (Fishilevich et al., 2016; Reinders et al., 2023; Rodrigues et al., 2021). A major focus of previous studies has been the identification of effective target genes, namely those that cause the highest mortality upon delivery of minimal doses of complementary dsRNA (Baum et al., 2007; Cedden and Bucher, 2025; Mehlhorn et al., 2021). A genome-wide RNAi screen in the red flour beetle *Tribolium castaneum* provided a comprehensive list of such effective target genes (Buer et al., 2025; Cedden and Bucher, 2025). These target genes were mainly involved in basic cellular functions, such as the proteasome pathway, which mediates the recycling of damaged or unnecessary proteins, and many of these targets showed cross-species efficacy (Buer et al., 2025; Cedden et al., 2025a). The findings from this screen aligned with efforts in target gene discovery in other pest species, suggesting that further improvements in pest control through target gene optimization may have reached a saturation point (Cedden and Bucher, 2025). Further improvements, however, may be achieved by optimizing the dsRNA sequence itself, which requires in-depth consideration of the underlying RNAi mechanism.

RNAi is initiated by dsRNA and enables sequence-specific gene silencing (Fire et al., 1998). Exogenously delivered dsRNA enters insect cells primarily through endocytosis and must reach the cytosol to engage the RNAi machinery (Koo and Palli, 2024; Zhu and Palli, 2020). In the cytosol, Dicer-2 processes dsRNA into siRNA duplexes. Each strand is typically approximately 21 nucleotides (nt) long in insects, although siRNA length can vary among species (Cedden et al., 2024; Cedden and Güney, 2026; Santos et al., 2019). A canonical 21-nt duplex contains a 19-base-pair (bp) region and 2-nt 3′ overhangs. Dicer-2 can cleave dsRNA processively, entering from a dsRNA end and generating successive siRNAs in approximately 21-bp increments (Cenik, 2011; Naganuma et al., 2021).

One strand of the siRNA duplex is retained as the guide strand by Argonaute-2 (Ago2), forming the RNA-induced silencing complex (RISC) and enabling cleavage of highly complementary RNA transcripts (Iwakawa and Tomari, 2022; Kawamata and Tomari, 2010; Sarkar et al., 2026). Guide-strand selection is influenced by duplex thermodynamic asymmetry, which favors retention of the strand with the more weakly paired 5′ end, i.e., with more A–U than G–C pairing (Lisowiec-Wąchnicka et al., 2019; Reynolds et al., 2004; Tomari et al., 2004). However, the relative contributions of individual terminal positions to strand bias remain poorly characterized in insect pests, limiting the use of this information in dsRNA design.

A recent study in *T. castaneum* identified siRNA sequence features that predict dsRNA regions with high insecticidal efficacy and incorporated these features into the dsRIP web platform (https://dsrip.uni-goettingen.de) (Cedden et al., 2025b). The dsRIP design algorithm averages predicted siRNA efficacy scores when searching for an optimal dsRNA target region, effectively assuming equal contributions from the possible siRNAs. However, different dsRNA sequences generate distinct siRNA abundance profiles (Cedden et al., 2024; Rodrigues et al., 2021). The RISC-bound siRNA pool generated by a given dsRNA may be shaped by several steps, including Dicer-2 processing (Fig. 1A), guide-strand selection (Fig. 1B), and post-loading modification and turnover (Fig. 1C). Their relative contributions *in vivo* remain unclear. Antisense guide strands can direct cleavage of the intended target mRNA, whereas sense guide strands do not target the corresponding site and may instead act on unintended transcripts (Fig. 1D). Therefore, understanding how dsRNA sequence shapes RISC-bound siRNA composition may support more effective dsRNA design and more informative off-target assessments.

**Figure 1.**
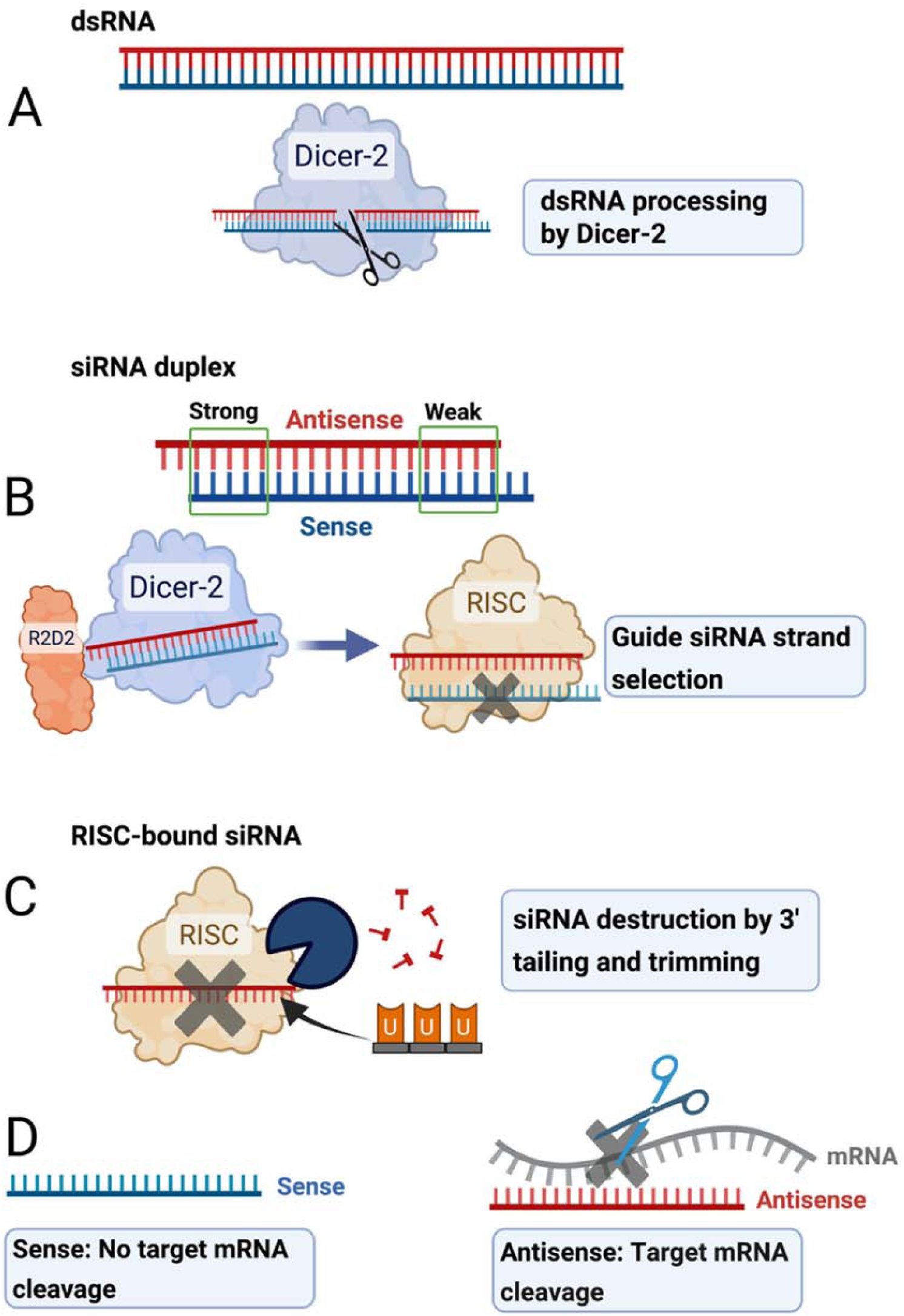
Potential steps shaping the RISC-bound siRNA pool after dsRNA delivery. (A) Dicer-2 processing generates the initial siRNA duplex pool. (B) Duplex thermodynamic asymmetry can bias guide-strand selection during RISC loading, involving Dicer-2 and R2D2. (C) Potential post RISC-loading modification and turnover mechanisms may alter the RISC-bound siRNA pool. (D) Antisense guide strands can direct cleavage at the intended target mRNA site, whereas the corresponding sense strands cannot, a distinction that affects the efficacy of gene silencing.

Following incorporation into RISC, small silencing RNAs can undergo 3′-end modifications that influence their stability and turnover (Iwakawa and Tomari, 2022). These include tailing, the addition of non-templated nucleotides, and trimming, the removal of nucleotides from the 3′ end. In *Drosophila*, HEN1-mediated 2′-O-methylation protects Ago2-associated siRNAs from these modifications (Ameres et al., 2010; Horwich et al., 2007). HEN1-family functions in small RNA stability have also been studied in *Caenorhabditis elegans* (Kamminga et al., 2012). Incomplete protection or structural changes associated with target recognition may expose small RNA 3′ ends to nucleotidyl transferases and exonucleases, promoting tailing and trimming (Ameres et al., 2010; Han and Mendell, 2023; Pisacane and Halic, 2017). The relevance of these processes to exogenously delivered insecticidal dsRNA remains poorly understood. By influencing the composition and persistence of the RISC-bound siRNA pool, terminal modifications may provide an additional layer of regulation affecting RNAi activity in pest insects.

Previous small RNA-sequencing (sRNA-seq) studies of siRNA abundance following dsRNA delivery have often examined only one or two dsRNA sequences and used total RNA extraction to recover dsRNA-derived small RNAs (Cedden et al., 2024; Guan et al., 2018). Limited sequence sampling restricts generalization, whereas total RNA preparations can include non-RISC-loaded siRNAs and mRNA cleavage products (Cedden, 2026). To address these limitations, we generated a RISC-bound sRNA-seq dataset following injection of 34 insecticidal dsRNAs targeting 11 essential genes in *T. castaneum* larvae. We examined associations of terminal sequence asymmetry, target region, and predicted self-folding with the antisense fraction of the RISC-bound siRNA pool. We then characterized siRNA tailing and putative trimming to explore whether terminal modification patterns were consistent with differential post RISC-loading siRNA retention.

## 2. Materials and Methods

### 2.1. Insects

The *T. castaneum* laboratory colony was maintained at 28 °C and 40% relative humidity under a 16 h light:8 h dark cycle in whole-wheat flour supplemented with 5% yeast powder.

### 2.2. dsRNA design

The dsRIP web platform (https://dsrip.uni-goettingen.de, v1.0) was used to design two to four dsRNAs for each of 11 essential target genes selected from Buer et al. (2025). The dsRNAs differed in target region, primarily the open reading frame (ORF) or 3′ untranslated region (UTR); one dsRNA targeted the 5′ UTR and was excluded from ORF-versus-3′ UTR comparisons. They also differed in predicted dsRIP scores, as described previously (Cedden et al., 2025b). Their lengths ranged from 298 to 347 bp, with a mean of 315.5 bp. Primer pairs (Supplementary Table 1) were designed using dsRIP and supplied by Integrated DNA Technologies (Germany).

### 2.3. dsRNA synthesis and injection

DNA templates for in vitro transcription were generated by overhang PCR using primers containing 5′ T7 promoter sequences. PCRs (50 µL) were performed with Phusion High-Fidelity DNA Polymerase (NEB, Germany) using 28–30 cycles and annealing temperatures of 58–60 °C. Amplicon lengths were verified by agarose gel electrophoresis, and the products were purified using the Gel and PCR Clean-up Kit (Macherey– Nagel, Germany).

DNA templates containing T7 promoters were transcribed using the MEGAscript T7 Transcription Kit (Thermo Fisher Scientific, Germany). The resulting dsRNAs were purified by lithium chloride precipitation according to the manufacturer’s instructions. The dsRNAs were denatured at 94 °C for 5 min and annealed at room temperature for 30 min. Annealed dsRNAs were verified on a 1.5% agarose gel.

Five late fifth-instar *T. castaneum* larvae were each injected with approximately 1 µL of injection buffer (5 mM KCl, 0.1 mM KH_2_PO_4_, 0.1 mM Na_2_HPO_4_, pH 6.8) containing 200 ng/µL dsRNA. Larvae were pooled and frozen in liquid nitrogen 3 days post-injection. A total of 41 samples were prepared: 38 from the 34 insecticidal dsRNA treatments (n = 1 or 2) and three from the dsmGFP control (n = 3).

### 2.4. RISC-bound small RNA sequencing

RISC complexes were isolated using the TraPR Small RNA Isolation Kit (Lexogen GmbH, Vienna, Austria) according to the manufacturer’s protocol (Grentzinger et al., 2020). Samples were crushed with pestles in 300 µL lysis buffer and homogenized by vortexing. Lysates were clarified by centrifugation at 10,000 × g for 5 min at 4 °C, loaded onto TraPR columns, mixed with the resin, and eluted by centrifugation. The elution step was repeated twice with 250 µL elution buffer, yielding a total of 750 µL RISC-containing eluate.

RISC-bound small RNAs were extracted with acid phenol:chloroform:isoamyl alcohol (25:24:1, pH 4.3; ROTI Aqua, Carl Roth) and precipitated with 0.3 M sodium acetate (pH 5.2) and isopropanol. Pellets were washed three times with ice-cold 80% ethanol and resuspended in 10 µL RNA elution buffer. Libraries were prepared using the NEBNext Multiplex Small RNA Library Prep Kit (NEB, E7560S), with 6 µL RISC-bound small RNA eluate as input. Final PCR amplification comprised 11 cycles. Amplified DNA was purified using the Monarch PCR & DNA Cleanup Kit (NEB), with a 7:1 binding-buffer-to-sample ratio, and eluted in 27.5 µL nuclease-free water. Library quality was assessed using an Agilent 2100 Bioanalyzer before and after selection of 135–150-bp cDNA fragments, corresponding to 15–30-nt small RNA inserts. Libraries were pooled and sequenced on a DNBSEQ-G400 platform (BGI, Hong Kong).

Raw sRNA-seq reads were adapter-trimmed and quality-filtered using Trim Galore v0.6, retaining reads with Phred quality >30. Cleaned reads were mapped to the respective dsRNA reference sequences using Bowtie v1.3, allowing up to three mismatches and reporting all valid mappings in both orientations. Small RNAs from the dsmGFP-injected control group did not map to the insecticidal dsRNA sequences. Perfectly matching 21-nt reads were extracted for the antisense-fraction analysis using samtools v1.2 and Python 3. Reads of additional lengths were examined separately for the mapping and terminal-modification analyses described below.

### 2.5. Analysis of siRNA features

For the four dsRNA treatments with two biological replicates, normalized siRNA abundances were averaged across replicates before calculating antisense fractions. For each siRNA pair, the antisense fraction was calculated as A/(A + S), where A and S denote the normalized abundances of antisense and sense strands, respectively. Only perfectly matching 21-nt reads were used. Sense and antisense features were paired computationally using the expected 19-bp paired region and 2-nt 3′ overhangs. Pairs were retained when at least one strand had non-zero abundance, yielding a total of 7,879 siRNA pairs for downstream analyses. Thus, the analyzed siRNA pairs represent computationally inferred precursor duplexes rather than intact duplexes isolated from RISC.

To investigate which nucleotide positions contribute to thermodynamic asymmetry, we compared nucleotides at the 3′ overhangs (positions −2 and −1) and paired terminal positions (1–6) with their respective symmetric counterparts at the opposite end of the siRNA pair (see Fig. 3C). Paired positions 1–6 were numbered inward from the terminal base pair at each duplex end. A value of +1 was assigned when the antisense 5′-side nucleotide was weakly paired (A or U) while the corresponding sense 5′-side nucleotide was strongly paired (G or C). A value of −1 was assigned for the converse configuration, with a strongly paired antisense 5′-side nucleotide (G or C) and a weakly paired sense 5′-side nucleotide (A or U). A value of 0 indicated that both ends had the same pairing strength. This scoring system quantified the bias toward weakly paired antisense 5′ ends, which is expected to favor antisense-strand selection during RISC loading. For each siRNA pair, the calculated ΔGC at each nucleotide position was correlated with strand-bias outcomes, with strand preference coded as 1 for antisense bias and 0 for sense bias. Pearson’s correlation coefficient, equivalent to a point-biserial correlation given the binary coding, was used to evaluate the relationship between ΔGC and strand bias.

The siRNA pairs were grouped by parameters calculated using dsRIP (https://dsrip.uni-goettingen.de, v1.2), including thermodynamic asymmetry score, GC%, lack-of-self-folding score, mRNA accessibility score, and target region (ORF or 3′ UTR). For the updated asymmetry score, positional ΔGC values at paired positions 1–5 were weighted by their correlations with strand bias and summed. The resulting scores were normalized to a 0–100 scale, with the highest score representing AU-rich pairing near the antisense 5′ end and GC-rich pairing near the sense 5′ end. Higher lack-of-self-folding and accessibility scores indicate less predicted antisense self-folding and greater predicted target mRNA accessibility, respectively. Multiple-group comparisons used the Kruskal–Wallis test followed by Dunn’s multiple-comparisons test. The ORF versus 3′ UTR comparison used the Mann–Whitney test.

### 2.6. Analysis of siRNA trimming and tailing

Non-templated terminal additions were characterized by nucleotide identity and length (1–3 nt), using 5′ additions as a directional control for 3′-end enrichment. Reads were classified by an exact templated siRNA core and terminal nucleotides that did not match the corresponding reference, rather than by total alignment mismatch count alone. For a 21-nt core, 22–24-nt reads were classified as tailed when the core matched perfectly and the 3′ or 5′ extension comprised one to three non-templated nucleotides.

For strand-specific analysis, we retained 3′ U, UU, or UUU extensions on either an exact 21-nt core or a shortened, exact 16–20-nt core. The latter were treated as putatively trimmed and U-tailed siRNAs. Each read identifier was counted once per siRNA feature to avoid duplicate counting of multiple alignments to the same feature. Relative U-tailing abundance was calculated separately for each strand as the combined count of U-tailed 21-nt cores and U-tailed 16–20-nt cores divided by the raw count of the corresponding perfectly matching 21-nt siRNA. For treatments with two biological replicates, relative U-tailing abundance was calculated separately for each replicate and then averaged. This measure is a relative read-abundance ratio, not a kinetic rate or a bounded fraction of modified molecules, and does not quantify untailed trimming products.

Strand comparisons included only siRNA pairs with non-zero normalized abundance and non-zero perfect 21-nt raw read counts for both strands; zero modification counts were retained, yielding a total of 4,291 siRNA pairs. Relative U-tailing abundances were compared across antisense GC-content, normalized lack-of-self-folding, normalized accessibility, and transcript-region categories. Multiple-group comparisons used the Kruskal–Wallis test followed by Dunn’s multiple-comparisons test.

## 3. Results

### 3.1. RISC-bound siRNA profiles generated from exogenous dsRNA

We generated a RISC-bound siRNA dataset from 34 insecticidal dsRNAs targeting 11 essential genes, with two to four dsRNAs analyzed per gene. After read filtering and computational pairing of complementary 21-nt features with 2-nt 3′ overhangs, 7,879 siRNA pairs had detectable abundance for at least one strand. We used these pairs to examine associations between siRNA features and antisense fraction, defined as antisense abundance divided by the combined abundance of antisense and sense strands.

Across the siRNA pairs, the mean antisense fraction was 0.521 and the median was 0.563 (95% CI of the mean, 0.511–0.530; Fig. 2A,B). These summaries indicate a slight overall antisense bias. Individual pairs nevertheless showed substantial heterogeneity, with fractions spanning 0 to 1 (Fig. 2A). Mean antisense fractions by target gene ranged approximately from 0.44 to 0.59 (Fig. 2C). Thus, gene-level averages were relatively close to equal antisense and sense abundance despite variation among individual siRNA pairs.

**Figure 2.**
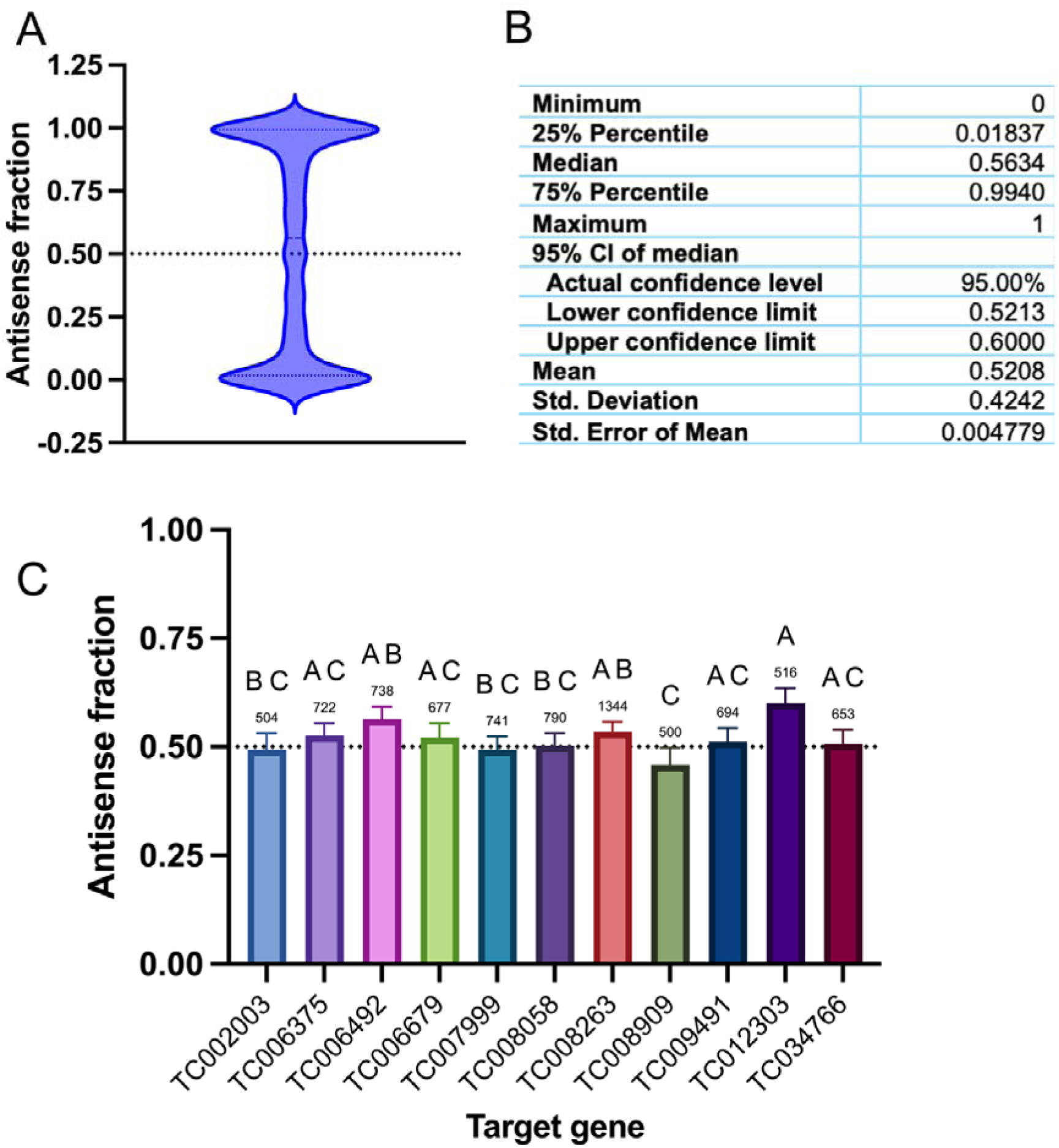
Distribution of antisense fractions in the RISC-bound siRNA dataset. (A) Antisense fractions across 7,879 siRNA pairs. The dotted reference line marks equal antisense and sense abundance (antisense fraction = 0.5). (B) Descriptive statistics, including the 95% CI of the median and the standard error of the mean. (C) Antisense fractions grouped by target gene. Bars show means ± 95% CIs; numbers indicate the number of siRNA pairs per gene. Groups were compared using the Kruskal–Wallis test followed by Dunn’s multiple-comparisons test. Groups sharing no letter differ at *P* < 0.05.

**Figure 3.**
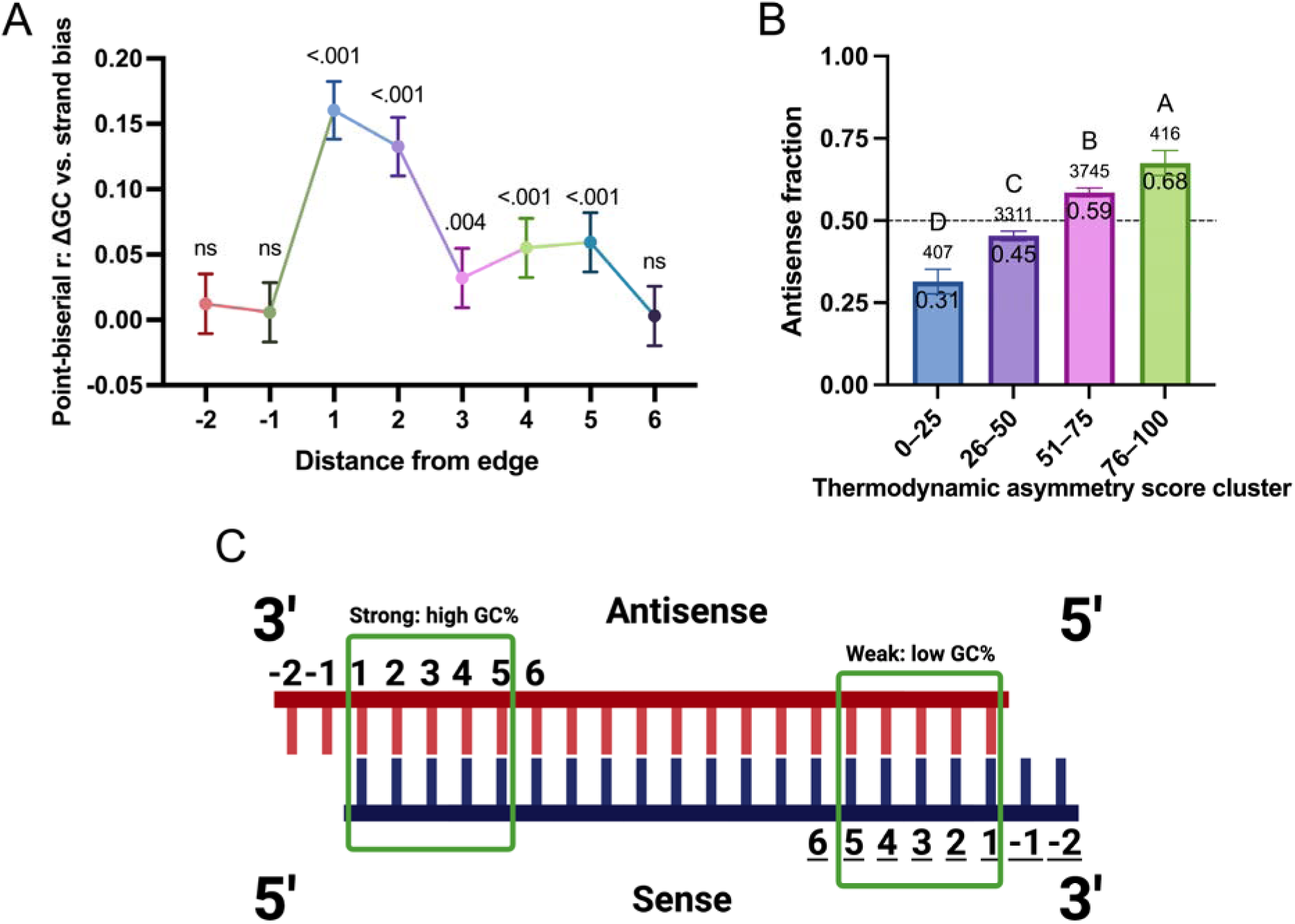
Association of terminal sequence asymmetry with siRNA strand bias. (A) Point-biserial correlations between positional ΔGC (3′ vs. 5′ end of the antisense strand) at unpaired overhang positions (−2 and −1) or paired positions (1– 6) and binary strand bias, defined by positive versus negative antisense-minus-sense abundance, across 7,879 siRNA pairs. P values are shown above the points; ns, not significant. (B) Antisense fractions grouped by normalized asymmetry score. Bars show means with 95% CIs; numbers indicate the number of siRNA pairs. Groups were compared using the Kruskal–Wallis test followed by Dunn’s multiple-comparisons test. Groups that do not share a letter differ at P < 0.05. (C) Schematic defining the terminal positions; paired positions 1–5 contribute to the updated score.

### 3.2. Terminal paired positions 1–5 correlate with siRNA strand bias

We used the antisense fraction in RISC to examine sequence features associated with guide-strand selection (Fig. 1B). Because the measurements were obtained 3 days after dsRNA injection, the observed antisense fractions are expected to reflect both initial guide-strand selection and subsequent differences due to siRNA turnover.

To resolve the contributions of individual terminal positions, we calculated positional ΔGC scores as proxies for local pairing strength asymmetry (Fig. 3C). Point-biserial correlations between ΔGC and binary strand bias were significant at paired positions 1–5 (*P* < 0.05, *r* = 0.03–0.16; Fig. 3A). Overall, AU-rich pairing near the antisense 5′ end and GC-rich pairing near the sense 5′ end were associated with antisense bias.

Positions 1 and 2 showed the strongest correlations (*r* = 0.16 and 0.13, respectively; both *P* < 0.001). Neither the two 3′-overhang positions nor paired position 6 showed a significant association. These results are consistent with a contribution of terminal pairing asymmetry to guide-strand selection, although the individual correlations were modest.

Using these positional correlations as weights, we calculated the updated asymmetry score described in Section 2.5 and grouped siRNA pairs into four score clusters. The same dataset was used to derive and examine the score, so the following comparisons describe its within-dataset association with antisense fraction rather than independent predictive validation.

The highest-score cluster (76–100%, *n* = 416) had a mean antisense fraction of 0.68, compared with 0.31 in the lowest-score cluster (0–25%, *n* = 407; Fig. 3B). These extreme clusters contained fewer siRNA pairs than the two intermediate clusters (*n* = 3,311 and 3,745), whose mean antisense fractions differed by 0.14. Thus, the score separated groups with different mean antisense fraction within this dataset. The increasing trend was also reported at the dsRNA-construct and gene levels and after omission of individual constructs or genes (Fig. S1).

### 3.3. Additional siRNA features are associated with antisense fraction

We next examined associations between antisense fraction and GC content, predicted self-folding, predicted target accessibility, and transcript region. These features may influence target interactions or siRNA persistence, but associations with final strand abundance do not by themselves establish when in the pathway they act.

Antisense fractions differed among GC-content clusters, with intermediate GC% associated with higher fractions than the lowest GC% category (*P* < 0.05; Fig. 4A). The cluster with the least predicted antisense self-folding had the highest mean antisense fraction (Fig. 4B). This pattern is consistent with, but does not directly demonstrate, preferential retention of antisense strands that can more readily interact with target transcripts.

**Figure 4.**
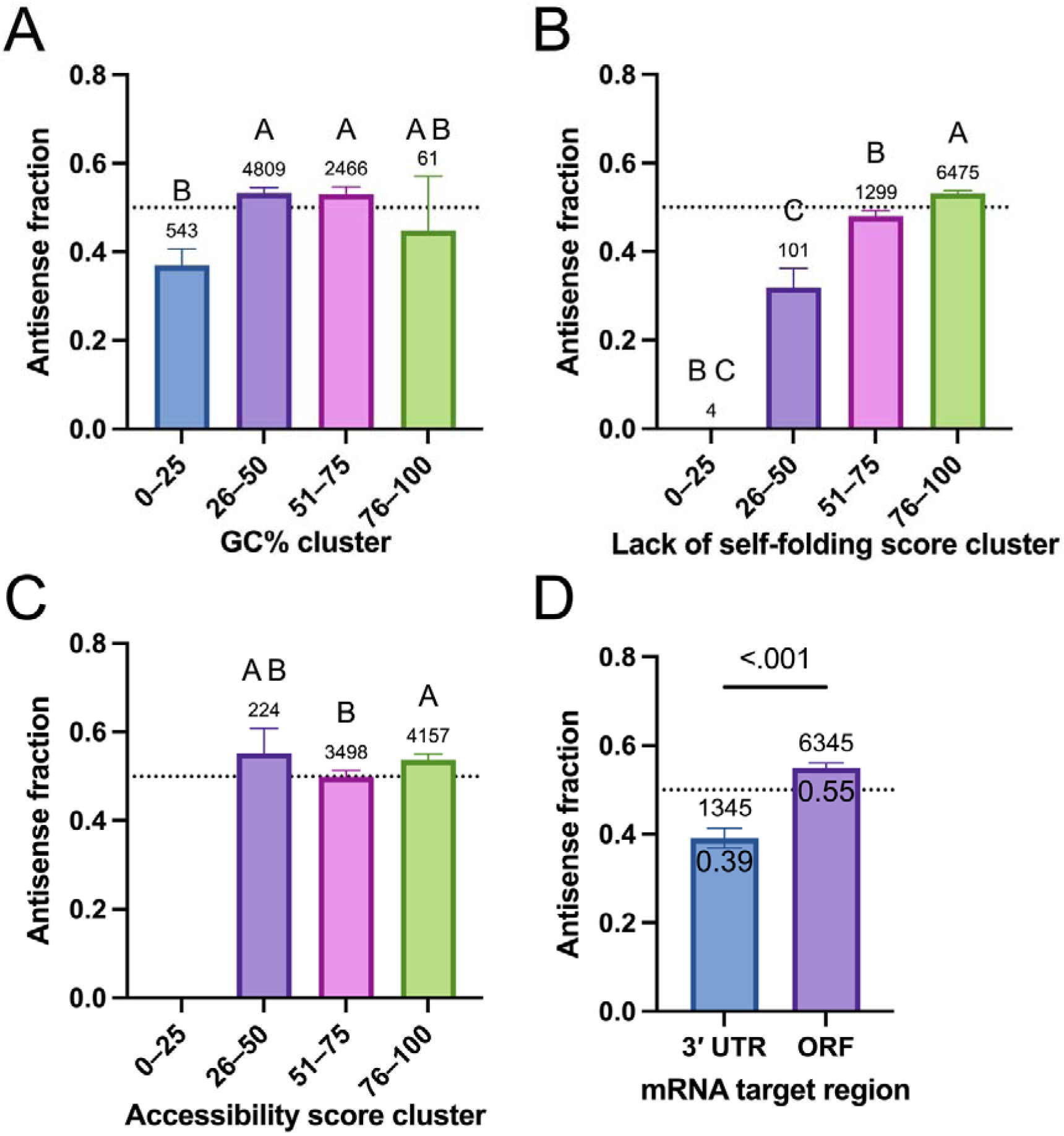
Associations between siRNA features and antisense fraction. siRNA pairs were grouped by (A) GC%, (B) normalized lack-of-self-folding score, (C) normalized mRNA accessibility score, or (D) target region. The dotted line marks equal antisense and sense abundance. Bars show means ± 95% CIs; numbers indicate the number of siRNA pairs. Panels A–C were analyzed using the Kruskal–Wallis test followed by Dunn’s multiple-comparisons test; groups sharing no letter differ at *P* < 0.05. Panel D was analyzed using the Mann–Whitney test.

Predicted mRNA accessibility was also associated with antisense fraction. The highest-accessibility cluster had a higher fraction than the adjacent intermediate cluster (*P* < 0.05; Fig. 4C), whereas the lower-accessibility cluster did not differ significantly from either group. The small number of siRNA pairs in low-score categories limits interpretation. ORF-associated siRNA pairs had a higher mean antisense fraction than 3′ UTR-associated pairs (*P* < 0.05; Fig. 4D).

Additional dsRNA-clustered and leave-one-construct-or leave-one-gene-out analyses of these features were consistent with these results (Fig. S2).

### 3.4. Terminal modifications in dsRNA-derived RISC-bound siRNAs

We next examined terminal-modification patterns that might accompany differences in post-loading siRNA persistence (Fig. 1C). We first characterized mapping mismatch classes and read lengths (Fig. 5A,B), then identified terminal additions using exact siRNA cores and non-templated extensions.

**Figure 5.**
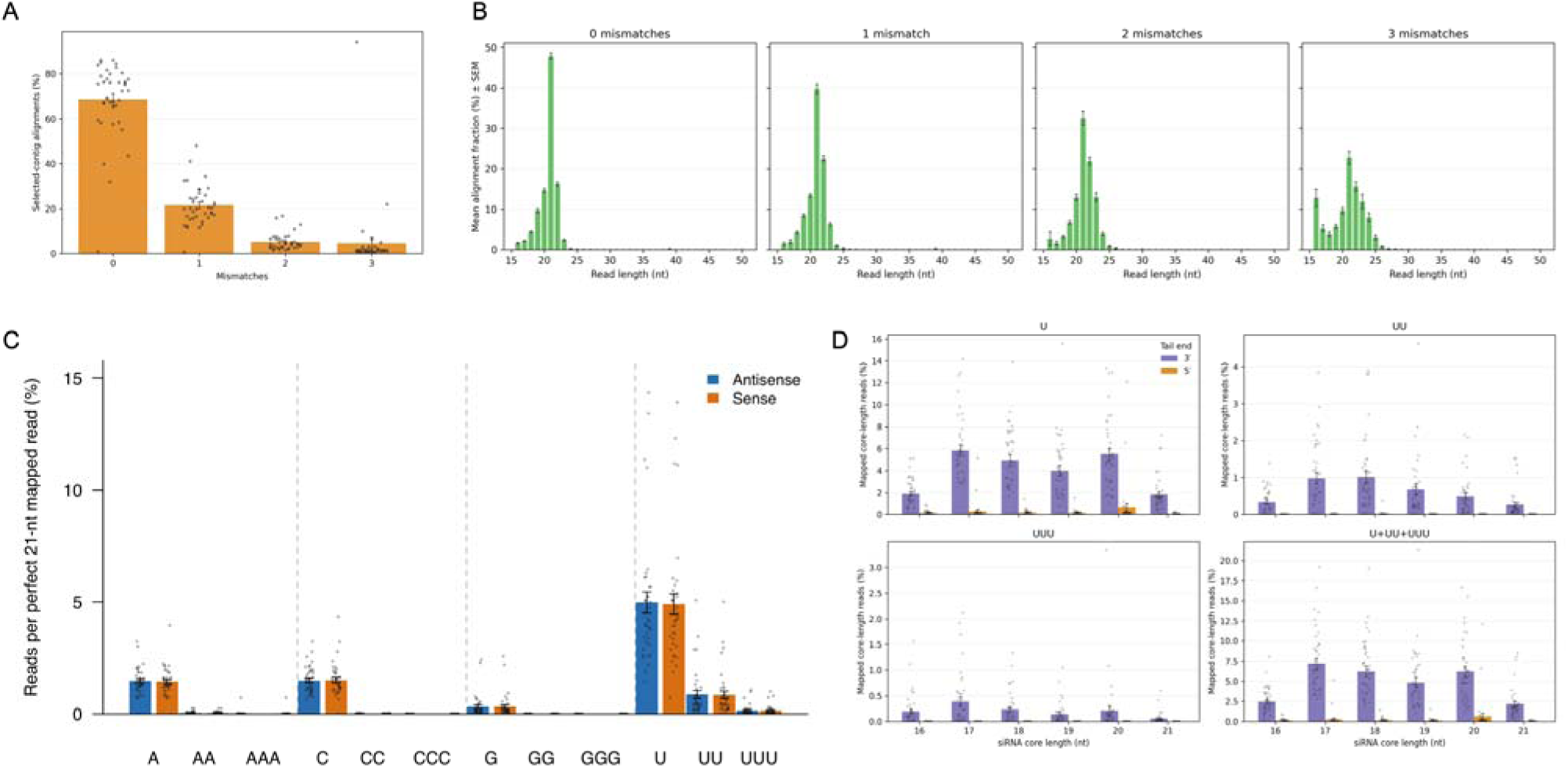
Mapping characteristics and non-templated tailing of dsRNA-derived siRNAs. (A) Percentage of alignments mapping to the reference contig assigned to each library, stratified by Bowtie mismatch class (0–3 mismatches). (B) Read-length distributions within these mismatch classes. (C) Abundance of 3′ homopolymer-tailed antisense and sense reads relative to perfect 21-nt mapped reads. (D) Abundance of U, UU, UUU, and combined U/UU/UUU tails across core lengths of 16–21 nt, comparing 3′ additions with a 5′-end directional control. In panels A, C, and D, bars show means ± SEMs and dots represent libraries.

A length of 21 nt predominated among perfectly matching reads, whereas reads containing one or more mismatches showed less precise length distributions (Fig. 5B), consistent with siRNA trimming and tailing in these mismatched reads. Among reads with a perfectly matching 21-nt core, non-templated 3′ tails were predominantly U additions, including up to three U extensions (Fig. 5C). Across the examined core lengths, 3′ U tailing was more abundant than the corresponding 5′-end control (Fig. 5D), supporting preferential 3′ modification, a known signature of small-RNA turnover.

### 3.5. ORF-associated sense siRNAs have higher relative U-tailing abundance

To relate terminal modifications to strand biases, we quantified reads with an exact 21-nt core followed by a non-templated 3′ U, UU, or UUU tail, together with putatively trimmed 16–20-nt cores carrying these tails. Their combined abundance was expressed relative to the corresponding perfectly matching 21-nt siRNA for each strand.

In both intermediate GC-content clusters, sense strands had higher relative U-tailing abundance than antisense strands (*P* < 0.05; Fig. 6A). No significant strand difference was detected in either extreme GC-content cluster (*P* > 0.05). The intermediate GC-content clusters also had higher antisense fractions (Fig. 4A).

**Figure 6.**
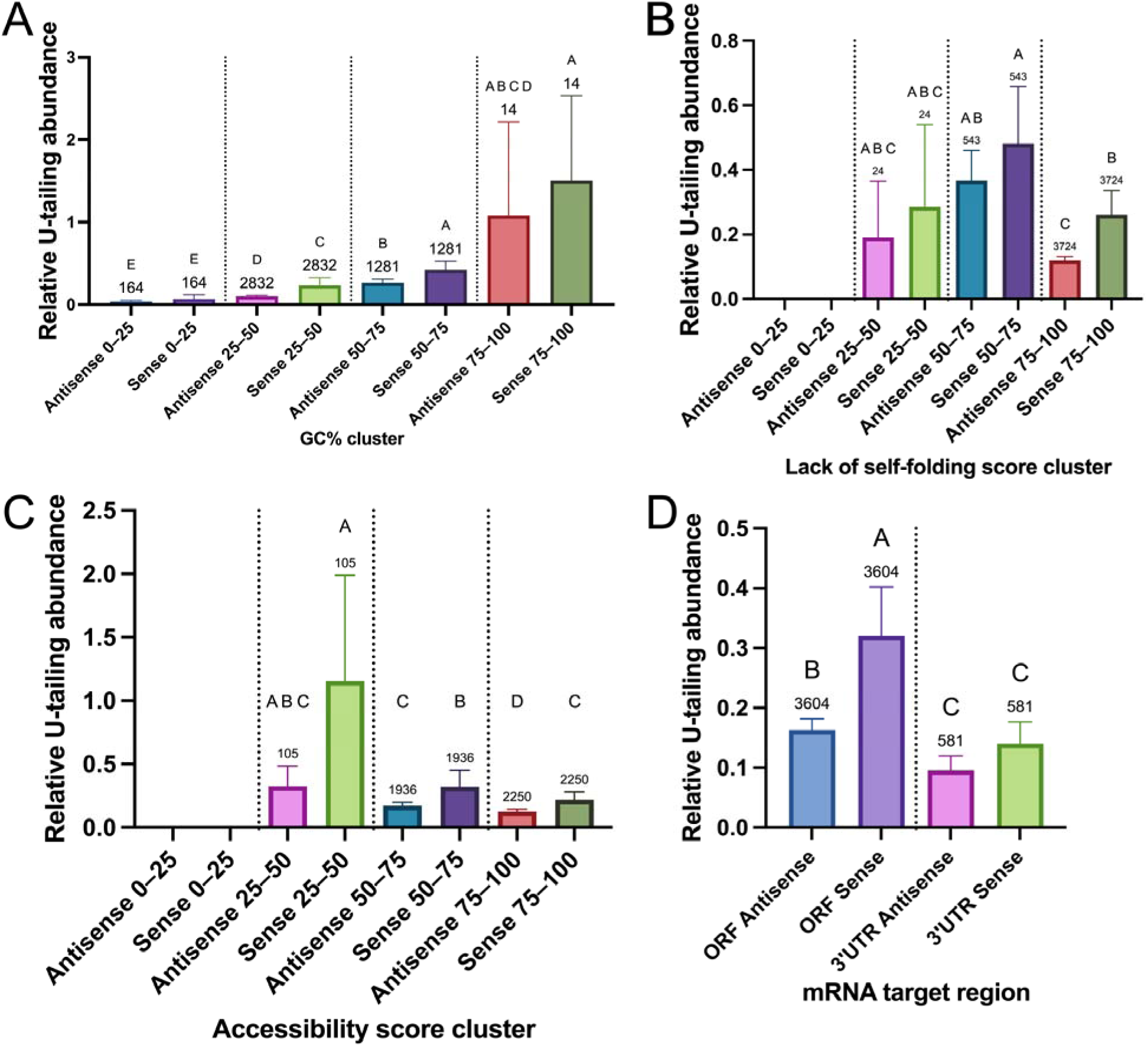
Strand-specific relative U-tailing abundance across siRNA feature groups. Relative U-tailing abundance was calculated as the combined counts of 3′ U-, UU-, or UUU-tailed reads with exact 21-nt or putatively trimmed 16–20-nt cores divided by the corresponding perfect 21-nt siRNA read count. Antisense and sense values are shown by (A) GC%, (B) normalized lack-of-self-folding score, (C) normalized mRNA accessibility score, and (D) target region. Bars show means ± 95% CIs; numbers indicate the number of siRNA pairs in each group. Groups were compared using the Kruskal–Wallis test followed by Dunn’s multiple-comparisons test; groups sharing no letter differ at *P* < 0.05.

Antisense strands in the cluster with the least predicted self-folding had the lowest mean relative U-tailing abundance (Fig. 6B). Within this cluster, sense strands had significantly higher relative U-tailing abundance than antisense strands (*P* < 0.05). This cluster also had the highest mean antisense fraction (Fig. 4B).

Antisense strands in the highest-accessibility cluster likewise showed low relative U-tailing abundance, including lower relative U-tailing abundance than the corresponding sense strands (*P* < 0.05; Fig. 6C).

Among ORF-associated pairs, sense strands had higher relative U-tailing abundance than antisense strands (*P* < 0.05; Fig. 6D). No significant strand difference was detected among 3′ UTR-associated pairs (*P* > 0.05). Relative U-tailing abundance was also higher for ORF-associated than for 3′ UTR-associated siRNAs within each strand category (*P* < 0.05).

These patterns are compatible with differential turnover of the two strands after RISC loading, additionally influenced by their features. However, the observed association does not establish preferential degradation of the sense strand or demonstrate its causal contribution to the observed antisense enrichment.

## 4. Discussion

RNAi offers a sequence-specific approach to pest control, but effective application requires suitable target genes and optimization of dsRNA sequences, in addition to suitable formulation strategies (Geisler et al., 2026; Kaarow et al., 2026). Genome-wide target screening in *T. castaneum* has provided a foundation for these efforts (Buer et al., 2025; Cedden and Bucher, 2025), and siRNA feature-based sequence optimization offers a complementary opportunity for improvement (Cedden et al., 2025b). Understanding how a given dsRNA sequence generates a mature RISC-bound siRNA pool in target and non-target insects may enable more efficacious pest control and improve the precision of off-target assessments (Chen et al., 2021; Chen and De Schutter, 2024; Mogren and Lundgren, 2017). Here, we identified associations between terminal pairing asymmetry, additional siRNA features, and antisense fraction in a RISC-bound siRNA dataset. We also observed strand-specific differences in putative uridylation and trimming patterns that were consistent with these associations.

Several limitations should guide interpretation of the current study. TraPR enriches RISC-associated small RNAs without establishing Ago2-specific occupancy (Grentzinger et al., 2020), and siRNA pairs were reconstructed computationally from separately sequenced strands. A single post-injection endpoint used in the study cannot by itself distinguish contributions of mechanisms at different steps (Fig. 1). For a complete picture, individual steps affecting RNAi efficacy, including non-specific RNase-mediated dsRNA degradation, initial siRNA duplex production by Dicer-2, RISC loading, and interactions between siRNAs and target mRNAs, should be investigated using high-throughput approaches (Cedden et al., 2025c; Guan et al., 2018; Peng et al., 2020).

Consistent with thermodynamic asymmetry models established in other systems (Lisowiec-Wąchnicka et al., 2019; Reynolds et al., 2004), sequence differences at terminal paired positions correlated with strand bias, with the correlation strength depending on the position. The updated weighted score, which includes 1-5 terminal paired positions, has been incorporated into dsRIP’s dsRNA efficacy prediction algorithm (v1.2) in place of the previous score derived from human-cell data (Cedden et al., 2025b). Its incremental predictive value requires testing on independent dsRNAs and through survival bioassays.

The strongest positional correlations occurred at the first two paired nucleotides. This is consistent with the established role of R2D2 in sensing duplex asymmetry during RISC loading in *Drosophila melanogaster* (Tomari et al., 2004; Yamaguchi et al., 2022). Preferential interaction with the more stable duplex end provides a mechanistic rationale for terminal sequence effects. However, R2D2 binding and its positional contributions were not measured directly in *T. castaneum* in this study.

Beyond terminal asymmetry, ORF targeting and reduced predicted antisense self-folding were associated with higher antisense fractions. ORF-associated siRNAs had a modest antisense bias, whereas 3′ UTR-associated siRNAs had a lower antisense fraction. This association is consistent with previous evidence favoring ORF-targeting dsRNAs (Cedden et al., 2025b; Rodrigues et al., 2021), but antisense fraction alone is not a measure of silencing efficacy. Independent efficacy tests are needed to determine whether the observed differences in antisense fraction improve target knockdown or mortality.

The association between transcript region and antisense fraction raises the possibility that interactions with target mRNAs influence the persistence of RISC-bound siRNAs (Ameres et al., 2010; Gajic et al., 2022; Nakanishi, 2016; Pisacane and Halic, 2017). In human cells, translating ribosomes can increase the accessibility of otherwise structured target sites and facilitate AGO2-mediated cleavage (Ruijtenberg et al., 2020). This mechanism provides a plausible context for differences between ORF- and UTR-associated siRNAs.

An antiviral context may also be relevant to the processing of exogenous dsRNA. The insect siRNA pathway is considered to be the primary defense against viruses that generate dsRNA during replication (Feng et al., 2026; Rawlings et al., 2011; Saleh et al., 2009). Most viral transcripts are composed largely of ORFs and UTRs are generally less accessible due to stable secondary structures (Slonchak et al., 2022). For these two reasons, we speculate that eliminating non-targeting sense strands while preferentially retaining complementary antisense strands targeting ORFs could enhance the efficiency of viral surveillance by RISC. In line with this hypothesis, a recent study demonstrated that virus-derived siRNA-loaded RISCs were predominantly associated with ribosomes to destroy polyadenylated viral transcripts rather than viral genomes (Silva et al., 2025). Interestingly, the strong association between RISC and ribosomes was observed regardless of viral infection status, supporting the idea that the results could be applicable to the processing of exogenously delivered dsRNA. These findings are consistent with the possibility that exogenous dsRNA-derived siRNAs preferentially interact with coding regions of target mRNAs during translation. This interaction could contribute to an increase in antisense fraction among siRNAs complementary to ORFs through preferential degradation of the sense strand, as suggested by the current study.

In *Drosophila*, 3′-terminal 2′-O-methylation protects Ago2-associated siRNAs from trimming and tailing (Ameres et al., 2010; Horwich et al., 2007). Differential methylation could therefore contribute to strand-specific modification, but methylation was not measured in the present study. Time-course measurements and perturbation of candidate modification enzymes would help distinguish altered loading, terminal modification, and degradation in future studies.

## 5. Conclusions

In conclusion, the current study suggests that the RISC-bound antisense fraction can be influenced by both guide-strand selection and subsequent interactions between RISC and the target mRNA, potentially through differential siRNA uridylation, putative trimming, and turnover. At the guide-strand selection step, base-paired nucleotide positions 1–5 at the duplex termini, especially the two terminal-most positions, were associated with antisense versus sense strand bias in RISC. Downstream of guide-strand selection, sense strands showed higher relative U-tailing abundance than antisense strands for siRNA pairs complementary to ORFs or more accessible regions, as well as in groups with reduced self-folding and intermediate GC content. Consistently, higher antisense fractions were observed under conditions associated with greater relative U-tailing abundance of the sense strand. Overall, these insights into factors associated with the composition of the RISC-bound siRNA pool may help improve the targeting of pest genes while minimizing potential off-target effects on non-target organisms.

## CRediT author contribution statement

Doga Cedden: Conceptualization, Methodology, Investigation, Formal analysis, Data curation, Visualization, Writing – original draft, Writing – review & editing, Supervision, Project administration, Funding acquisition. Gözde Güney: Investigation, Writing – review & editing. Paul Plathner: Validation, Writing – review & editing. Stefan Scholten: Methodology, Resources, Writing – review & editing.

## Acknowledgements

The authors thank Prof. Dr. Gregor Bucher for helpful discussions and for providing *T. castaneum* larvae. D.C. and G.G. were supported by scholarships from the German Academic Exchange Service (DAAD) Doctoral Programmes in Germany and by the German Research Foundation (DFG) through the Walter Benjamin Programme (Project No: 585515648 and 573363460). The graphical abstract and Figure 1 were created using BioRender.com without AI-generated images. Open Access funding was enabled and organized by Projekt DEAL.

## Data availability

The datasets supporting the conclusions of this article are provided as Supporting Datasets 1 and 2. The RISC-bound small RNA sequencing data have been deposited in the NCBI Sequence Read Archive under BioProject ID PRJNA1157760 (https://identifiers.org/ncbi/bioproject:PRJNA1157760)

