## Supporting information for "RISC-bound small RNA sequencing provides insights into guide strand selection and siRNA trimming and tailing following insecticidal dsRNA delivery"

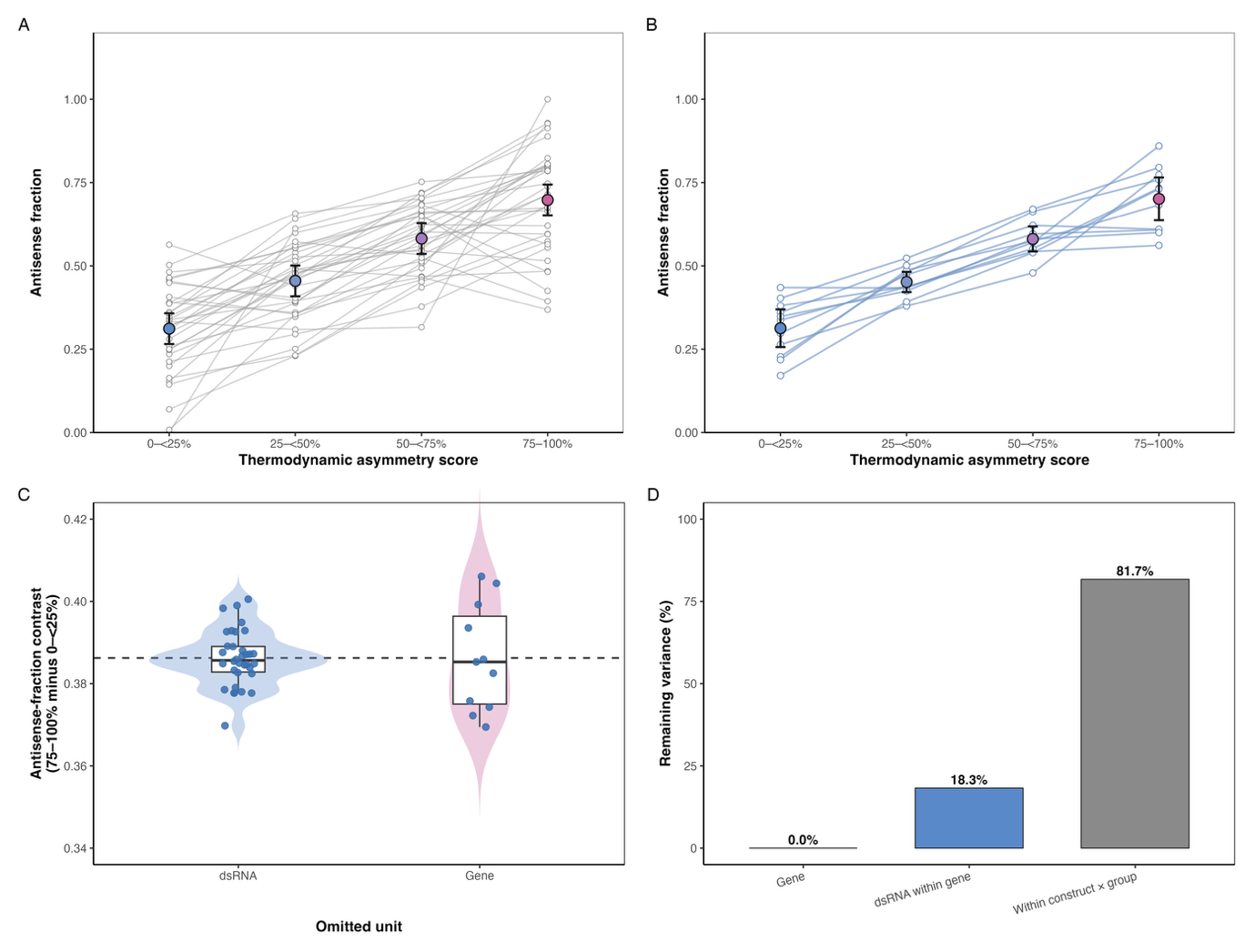

**Figure S1 Robustness of antisense enrichment across thermodynamic asymmetry-score clusters.** (A) Antisense fractions were averaged within each dsRNA construct (n = 34); lines represent constructs and large markers show construct-level mixed-model estimated means with 95% confidence intervals. (B) The same analysis after aggregation to equally weighted gene-level means (n = 11). (C) The contrast between the highest and lowest asymmetry groups after serial omission of one dsRNA construct or one gene. (D) Variance components after score group is included in the construct-level model. Individual siRNA rows were summarized within dsRNA constructs before inference; the score-group effect was significant (likelihood-ratio test, P < 0.001).

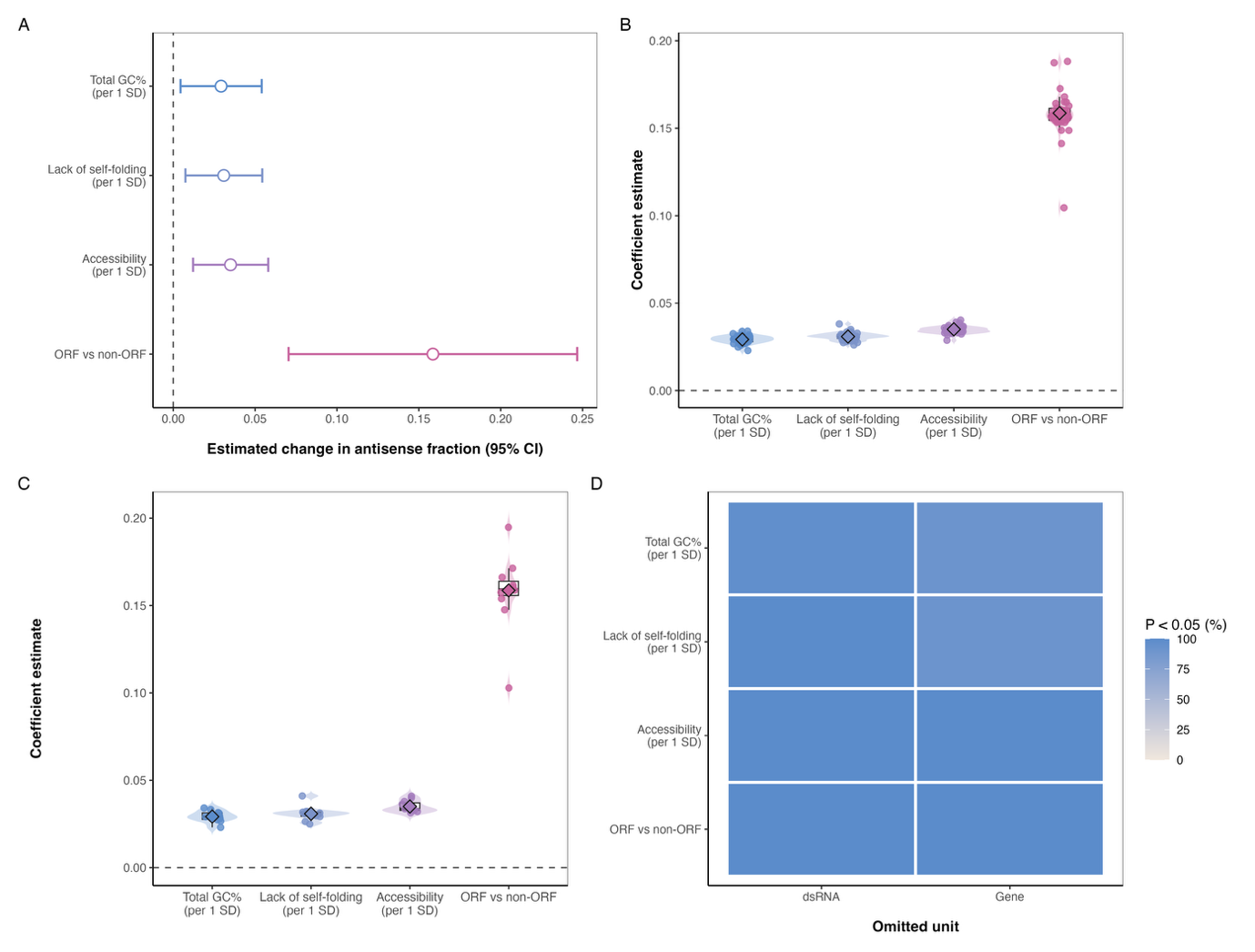

**Figure S2 Robustness of additional predictors of antisense fraction.** (A) Estimates and 95% confidence intervals from the multifactor model for total GC%, lack of self-folding, accessibility, and ORF status. (B,C) Coefficient distributions after serial omission of each dsRNA construct or gene, respectively; diamonds indicate the full-data estimate. (D) Percentage of leave-one-out fits retaining P < 0.05. Standard errors were clustered by dsRNA construct; continuous predictors are expressed per full-dataset standard deviation, and ORF is relative to non-ORF.

**Supplementary Table 1. Primer sequences used for in vitro transcription templates of 34 different essential-gene complementary dsRNAs**

| **Gene : dsRNA** | **Forward primer*** | **Reverse primer*** |
| --- | --- | --- |
| TC002003:dsRNA-1 | GCGAAGGAGGAATCAGTTGC | ATTTGTCGTTTTTAGGGACT |
| TC002003:dsRNA-2 | AAGCTCCCCACCCTTTGTTC | GCACGTCTGACCTTCGTAGT |
| TC002003:dsRNA-3 | ACTACGAAGGTCAGACGTGC | TCGCTGTACTCGTCCTCTGA |
| TC006375:dsRNA-1 | TCAGCACTTTCACATCGTACGA | TCCAATAACTGGGTGTATGCCA |
| TC006375:dsRNA-2 | TGCCGATCGAAAACCTGGAA | CCTCACGGACTTCCATCTCG |
| TC006375:dsRNA-3 | CGAAGCGTTTGGTGTCACAG | AAAGCTGTTTGCACAACAAA |
| TC006492:dsRNA-1 | ACCAGACAAACGTCAAATCA | GCCAACCCTGTGTCAGACTC |
| TC006492:dsRNA-2 | GACACCGACGACCCCAAATA | GGCTCAATCCCCAATTTGACG |
| TC006492:dsRNA-3 | ACGAAATTGACGCCATAGGC | GTGCACACGCTCCTGATCT |
| TC006679:dsRNA-1 | GGAACAGCCAATCCCGCTAA | TGTTGTTTGGTTGAAGTGGTCG |
| TC006679:dsRNA-2 | AAGCCGAACAGTGCACTCTT | GAACCTGGGGCACGACTAAA |
| TC006679:dsRNA-3 | CACCGAACTATGGGAAGCCA | CTCGCAAATCTGGACGCATG |
| TC006679:dsRNA-4 | ACGACTGACACTGACTTCACA | ACACAATTACTCCACTTCTTCT |
| TC007999:dsRNA-1 | TCATCGTAACATAACCCCAA | GAAACTGGCCGATGACAAGC |
| TC007999:dsRNA-2 | GCTGATACCCTCGACCCG | TCGTGCTCGCTTTCATCCTT |
| TC007999:dsRNA-3 | ACGATTGACCGGGAATTGCT | CCTTGGCCAGCATTGTCTTG |
| TC008058:dsRNA-1 | GGTGTGAGCAAGAAGACCCA | GGCACTTTGTCCGTACGAGA |
| TC008058:dsRNA-2 | TTCCTGCCGGTTGTACTGTC | TCCTTCGGCCCTCTTAAGAA |
| TC008058:dsRNA-3 | CCCTGCTGTTATGGTACGCT | TCAAAAGACCATCGCCGAGC |
| TC008263:dsRNA-1 | GCCAGTTACAGCCCAACCTA | CGTCCCGGTCTGAGTTCTTG |
| TC008263:dsRNA-2 | TGAGTACACTGTTCGCATTTCC | CCAAATAGAGTTTACCGATCGT |
| TC008263:dsRNA-3 | GACTCATTCCATGGACGCCA | TGTCGGAAATGCCAGTCCAA |
| TC008263:dsRNA-4 | GGGGGAAGAACCAGTCCAAT | TCCAACAATTGACAACAACAGA |
| TC008909:dsRNA-1 | CAGCAGCTACTTGCCTCTCT | AGCCTCTCATGGATGTTAACCA |
| TC008909:dsRNA-2 | CGAATGCGCCCAAAAACTCA | GGCCGAATACAGAGCCAACT |
| TC008909:dsRNA-3 | CGAATGTCGAAGAAGCAGCG | TGTCGTTCAGTGTGTCCAGG |
| TC009491:dsRNA-1 | ACTTGTACACAGTCATATTCGT | TGAAGTGCATACAAAATATCT |
| TC009491:dsRNA-2 | AAGAATTCGCTGTTTCCGGC | ACGAAATAACCCCAGCTGCT |
| TC009491:dsRNA-3 | TGTTTATGCTCGCCTCCAGC | CACTTCATCGCTGGCTTTGG |
| TC012303:dsRNA-1 | CGACTGTGTTAAATCAAGCCA | TCTCGTTTTGGGCCCTCTTC |
| TC012303:dsRNA-2 | CCTTTTCCAGCCTCCGACTC | AGTCGCTCAAGATGGAAAGCA |
| TC034766:dsRNA-1 | GAAGCCGAAAACTGCGACTG | CCAAATGGTTCAAGTCGCCG |
| TC034766:dsRNA-2 | GGGAGGTACTTGACAGTGGC | CCATCTCGTCCATTCCTTCCC |
| TC034766:dsRNA-3 | ACAACGAAGCCACAGTTGCT | TGAGTAAAGTGCCCATGCCG |
| dsmGFP control | GGACCCTGACCTACGGCCTATT | GGTGCCGTCCTCGTACTAGTTGATGC |

* The T7 promoter sequence “GAATTGTAATACGACTCACTATAGG” was added to the 5’ ends of both forward and reverse primers
